# A new approach for quantifying phosphorus requirement in an Amazonian fish using CT-scanning

**DOI:** 10.1101/2020.03.26.009886

**Authors:** Ludmila L.C. Menezes, Janaina G. A. Santos, Igo G. Guimarães, Delma M. C. Padua, Vânia M. V. Machado, Cristielle N. Souto

## Abstract

Phosphorus (P) is an essential mineral for fish growth, as it plays pivotal roles in skeletal development and energy transfer reactions. However, the dietary requirement of this mineral is variable among fish species and the growth stages. Thus, this study aimed to determine the digestible P (dP) requirement for tambaqui in the initial growth stage (± 17 to 150g) using growth data, mineralization of the whole body, vertebrae and scales, as well as blood chemistry as response parameters. A total of 192 tambaqui juveniles of approximately 17 ± 0.85 g were stocked into a water recirculation system. Fish were assigned to 24 70L-tanks using randomized block design (two floors) with six treatments (1.3, 2.4, 4.8, 6.3, 7.8 and 8.8 g kg^−1^ dP) and four replicates. Fish were fed six semipurified diets with increasing levels of dP for 90 days. The dietary requirement of P was estimated using regression models (P < 0.05). Duncan and SNK multiple range tests were used when regression models were not fitted. No mortality or apparent signs of P deficiency were observed. All performance variables were improved with increasing levels of dP in the diet. Based on weight gain, the P requirement was 6.3g kg^−1^ diet while for increased carcass mineral deposition was 6.6g kg^−1^ diet and for adequate mineralization of vertebrae the requirement was 4.75 g dP kg^−1^ diet. The blood chemistry parameters were greatly affected by the dietary P level, except for serum calcium. Thus, the dietary dP requirement for tambaqui juveniles in the early stage was 6.3 g kg^−1^ diet based on growth and bone mineralization.

## 1. Introduction

Phosphorus (P) is the first limiting nutrient for algae growth in freshwater ecosystem. Additionally, its association with nitrogen overload could induce the eutrophication of water bodies. Thus, P discharge by farmed fish might contribute to the pollution of water reservoirs and reduce the sustainability of the aquaculture production. In this context, Brazil is responsible for 6% of the worldwide phosphorus discharge to the aquatic environment [1] which poses a significant impact for the expected growth in aquaculture predicted for the following decades [2] once the soluble P removal from the water is complex and still inefficient [3]. Therefore, finding ways to reduce P discharges by aquaculture is a key point for the future of freshwater fish production [4] [3].

One of the suggested ways to efficiently reduce the P discharge from fish farming is by strictly adjusting the P intake to the specific requirement of a fish species in each developmental stage once during fish growth the nutrient requirement changes according to the physiological status and environmental changes. For instance, the P requirement tends to reduce according to the growth stage mainly due to the limitation of bone tissue growth as fish reaches the adult stage [5]. Thus, defining the minimum dietary P requirement in different growth stages is one of the first steps to reduce the P discharge in aquaculture production.

However, the P requirement in animals diverge depending on the response parameters evaluated and the models used to estimate the requirement [6]. For instance, P requirement based on weight gain are lower than those estimated based on bone mineralization [7–10]. The use of P requirement estimates based on WG have led to several bone deformities in salmonids imposing a threat to the welfare of fish and the quality of the final product [10–12]. This has been used as role for most of the aquacultured fish species, at least for Nile tilapia and carp aquaculture [8, 13]. However, limited data on other fish species are available and further extrapolations should be made with care.

Tambaqui *(Colossoma macropomum)* is the second most farmed species in Brazil and possess an important role on food security for Latin American countries [14]. Additionally, the high P discharge in tambaqui aquaculture might be a serious problem for the Amazonian aquatic ecosystem where most of the tambaqui farms are located. Therefore, the precise P formulation in tambaqui diets is of utmost importance for protecting this ecosystem.

Although the importance of this species for aquaculture, information on its nutrient requirements are limited and, to the best of our knowledge, just one study have reported the phosphorus requirement for tambaqui in the grow-out stage (~ 150g) [7]. Therefore, we designed a trial to determine the P requirement for tambaqui juveniles (from 15 to 150g) using different response parameters. Additionally, for the first time in a requirement study, we provide evidence that the P requirement for whole-body mineralization might be similar to the estimate based on weight gain while the requirement for bone mineralization are markedly low in fish with dense mineralized bones.

## 2. Materials and methods

All the procedures involving animals in this study was approved by the Animal Ethics Committee of the Universidade Federal de Goiás (protocol # 063/15).

### 2.1. Experimental diets

A basal semi-purified diet was formulated to meet the protein (360 g kg^−1^) and energy (14.02 Mj kg^−1^ diet) requirements for tambaqui and be deficient in phosphorus (1.5 g kg^−1^). Then, incremental levels of dietary digestible phosphorus were obtained by supplementing the basal diet with potassium phosphate to achieve the following levels: 3.0, 4.5, 6.0, 7.5 and 9.0 g kg^−1^ (Table 1). Digestible phosphorus content of potassium phosphate was calculated according to a previous study of our group [15].

**Table 1.**
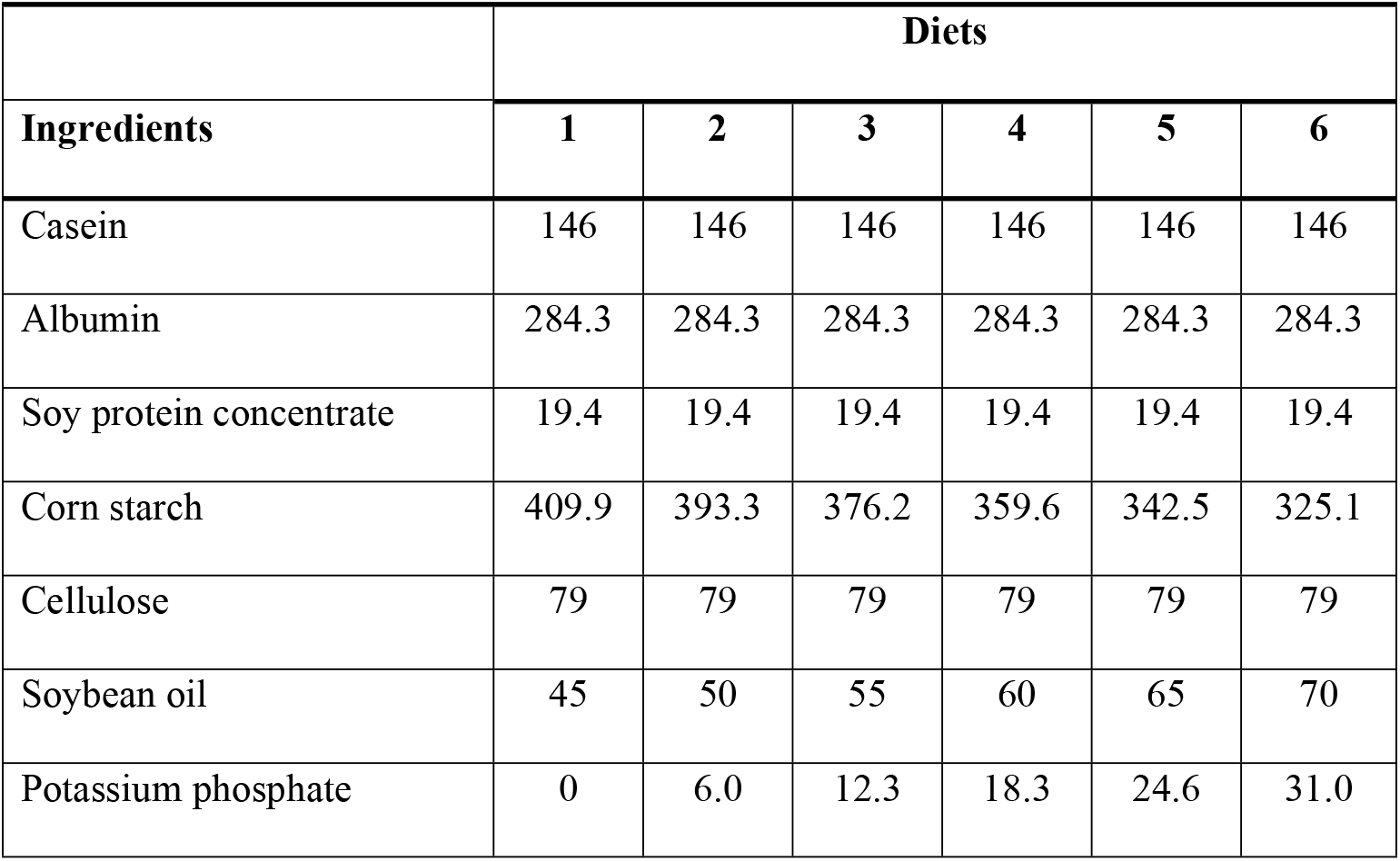
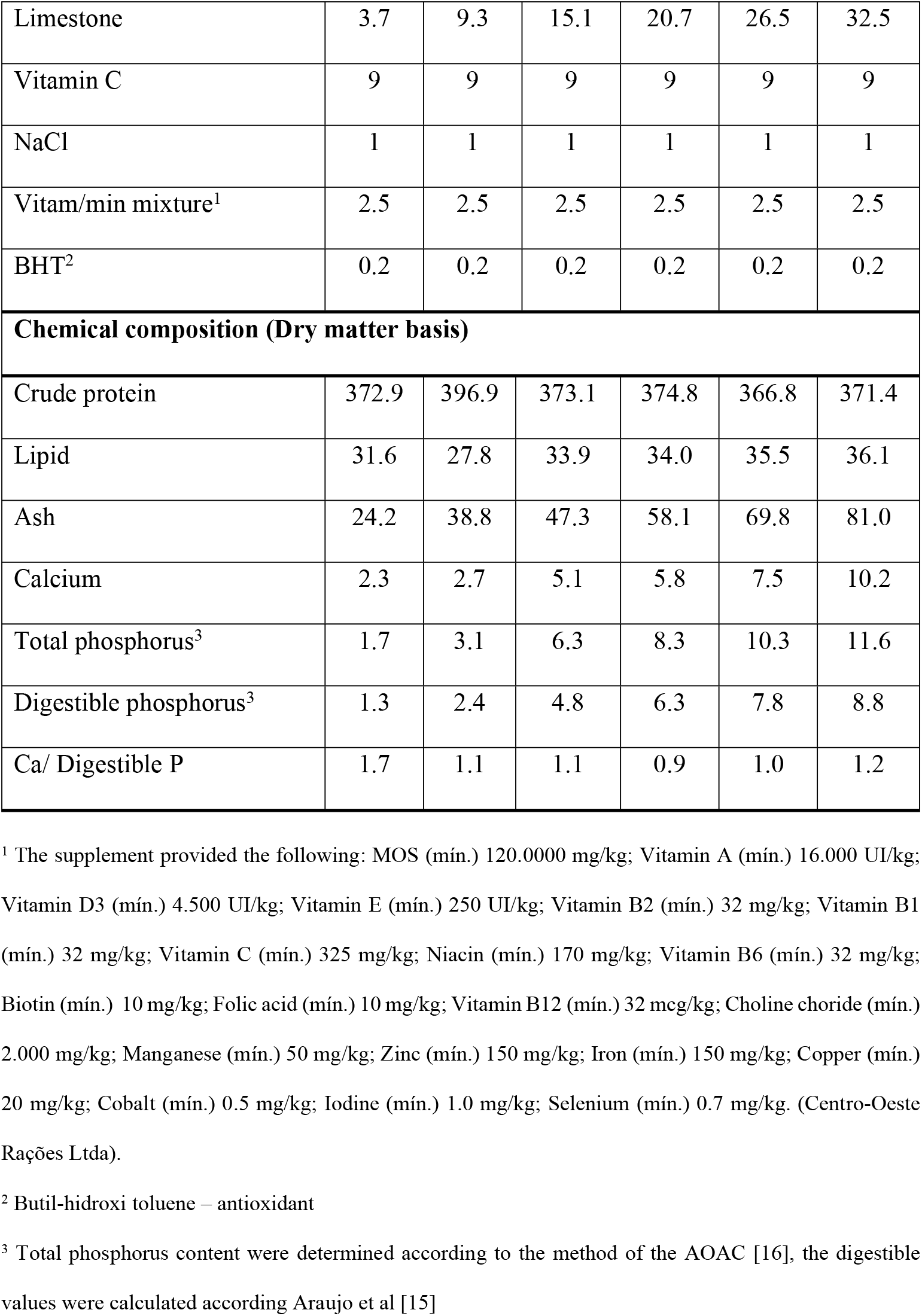
Ingredient and chemical composition of experimental diets (g kg^−1^)

|  | **Diets** |  |  |  |  |  |
| --- | --- | --- | --- | --- | --- | --- |
| **Ingredients** | **1** | **2** | **3** | **4** | **5** | **6** |
| Casein | 146 | 146 | 146 | 146 | 146 | 146 |
| Albumin | 284.3 | 284.3 | 284.3 | 284.3 | 284.3 | 284.3 |
| Soy protein concentrate | 19.4 | 19.4 | 19.4 | 19.4 | 19.4 | 19.4 |
| Corn starch | 409.9 | 393.3 | 376.2 | 359.6 | 342.5 | 325.1 |
| Cellulose | 79 | 79 | 79 | 79 | 79 | 79 |
| Soybean oil | 45 | 50 | 55 | 60 | 65 | 70 |
| Potassium phosphate | 0 | 6.0 | 12.3 | 18.3 | 24.6 | 31.0 |
| Limestone | 3.7 | 9.3 | 15.1 | 20.7 | 26.5 | 32.5 |
| Vitamin C | 9 | 9 | 9 | 9 | 9 | 9 |
| NaCl | 1 | 1 | 1 | 1 | 1 | 1 |
| Vitam/min mixture<sup>1</sup> | 2.5 | 2.5 | 2.5 | 2.5 | 2.5 | 2.5 |
| BHT<sup>2</sup> | 0.2 | 0.2 | 0.2 | 0.2 | 0.2 | 0.2 |
| **Chemical composition (Dry matter basis)** |  |  |  |  |  |  |
| Crude protein | 372.9 | 396.9 | 373.1 | 374.8 | 366.8 | 371.4 |
| Lipid | 31.6 | 27.8 | 33.9 | 34.0 | 35.5 | 36.1 |
| Ash | 24.2 | 38.8 | 47.3 | 58.1 | 69.8 | 81.0 |
| Calcium | 2.3 | 2.7 | 5.1 | 5.8 | 7.5 | 10.2 |
| Total phosphorus<sup>3</sup> | 1.7 | 3.1 | 6.3 | 8.3 | 10.3 | 11.6 |
| Digestible phosphorus<sup>3</sup> | 1.3 | 2.4 | 4.8 | 6.3 | 7.8 | 8.8 |
| Ca/ Digestible P | 1.7 | 1.1 | 1.1 | 0.9 | 1.0 | 1.2 |

<sup>1</sup> The supplement provided the following: MOS (mín.) 120.0000 mg/kg; Vitamin A (mín.) 16.000 UI/kg; Vitamin D3 (mín.) 4.500 UI/kg; Vitamin E (mín.) 250 UI/kg; Vitamin B2 (mín.) 32 mg/kg; Vitamin B1 (mín.) 32 mg/kg; Vitamin C (mín.) 325 mg/kg; Niacin (mín.) 170 mg/kg; Vitamin B6 (mín.) 32 mg/kg; Biotin (mín.) 10 mg/kg; Folic acid (mín.) 10 mg/kg; Vitamin B12 (mín.) 32 mcg/kg; Choline choride (mín.) 2.000 mg/kg; Manganese (mín.) 50 mg/kg; Zinc (mín.) 150 mg/kg; Iron (mín.) 150 mg/kg; Copper (mín.) 20 mg/kg; Cobalt (mín.) 0.5 mg/kg; Iodine (mín.) 1.0 mg/kg; Selenium (mín.) 0.7 mg/kg. (Centro-Oeste Rações Ltda).
<sup>2</sup> Butil-hidroxi toluene – antioxidant
<sup>3</sup> Total phosphorus content were determined according to the method of the AOAC [16], the digestible values were calculated according Araujo et al [15]

The ingredients were sieved to a mesh diameter less than 500 μm to adjust the DGM of each ingredient. Then, they were thoroughly mixed and soybean oil and water was added. The moisten mixture was pelleted in a meat grinder adapted to manufacture pelleted purified diets. The pellets were dried in an air forced oven at 55 °C for 24h, roughly grinded and sieved to present similar size to the open mouth of the fish. The diets were stored in airtight bags at −20 °C until the start of the experiment.

### 2.2. Experimental fish, facilities and design

Fish were acquired from a commercial fish farm (Aquicultura São Paulo). Firstly, the animals were stocked into three 1000L quarantine tanks and a protocol for disinfection was used to eliminate parasites. Fish were maintained during three weeks in these aquaria and fed a commercial fish diet. When the feeding of fish returned to normal, the basal diet was used for 10 days to acclimatize the fish to the experimental diets. 192 tambaquis with 17 ±0.5g were randomly assigned to 24 70L-aquaria connected to a closed recirculation water system.

A randomized block design with six treatments (dietary phosphorus levels) and four replicates was used. Since we had two sets of 12 aquaria in different floors (see the schematic view of the experimental settings), we decided to block each floor to control the differences between the two sets. Each experimental unit (aquarium) was stocked with eight fish, comprising 32 fish per treatment. Experimental diets were assigned to the tanks and fish were fed ad libitum two times a day (at 09:00 and 16:00).

Water quality parameters, such as temperature, pH, dissolved oxygen, alkalinity and total ammonia were monitored in a daily basis and were kept within the normal range for maximum growth and health of tambaqui [7].

Fish were anesthetized with eugenol (100mg/ 1L water) before all handling procedures and the animals used for tissue samplings were killed by using a megadose of the anesthetic [17].

### 2.3 Chemical analysis

Twenty fish were randomly sampled at the beginning of the trial for determining the initial whole-body chemical composition and for calculation of the nutrient retention. At the end of the trial (90 days), all fish were starved for 24h, weighed and tissues were sampled. Two fish per tank were sampled at the end of the trial for whole-body chemical composition.

Diets and fish samples were analysed in duplicates for chemical composition. Moisture was determined in an oven at 105 °C for 24h; protein (nitrogen * 6.25) was determined using microKjeldahl method; total lipid was determined using a Soxhlet extractor; and ash was determined by incineration in a furnace at 550 °C for 5 h [16].

Eight fish per treatment were dissected for scales and vertebrae samplings. Vertebrae were washed with deionized water to remove all remaining connective tissue, then lipids were removed by extraction in a chloroform to methanol solution (1:1) for 6h. The fat free vertebrae and scales were dried in an air oven at 65 °C for 24h, ground and prepared for acid digestion [18].

Mineral content of diets, whole-body, vertebrae and scales were determined after digestion with a nitropercloric solution (3:2) for 3 h at 300 °C. After digestion, the mineral extracts were diluted in 50 mL deionized water and minerals were determined. Phosphorus was determined using a colorimetric method while Ca, Mg, Mn and Zn were determined by atomic absorption spectrometry [16].

### 2.4 Growth performance parameters

Fish were weighed every 15 days and feed intake was recorded in a daily basis. Survival, final weight (FW), daily weight gain (DWG), feed intake (FI), feed conversion ratio (FCR), protein efficiency ratio (PER), protein retention (PR), and phosphorus utilization (PU) were evaluated according to the following equations:

Daily weight gain (DWG) = final weight (g) – initial weight (g) / length of the experiment

Feed intake (FI) = Feed consumption (g) / fish weight (g) / length of the experiment

Feed conversion ratio (FCR) = Dry feed fed (g) / wet weight gain (g)

Specific growth rate (SGR) = 100 x [(Ln Final Weight - Ln Initial Weight) / days of experiment]

Protein efficiency ratio (PER) = Wet weight gain (g) / protein intake (g)

Protein retention (PR) = [(final weight (g) x final whole-body protein content (g)) – (initial weight (g) x initial body proeint content (g)/ ingested protein (g)] x 100; Phosphorus utilization (PU) = Wet weight gain (g) / phosphorus intake (g)

### 2.5 Hematology

Three fish per tank were collected and bled by caudal puncture with 1 mL syringe rinsed with without anticoagulant for the serum biochemistry analysis.

The blood sample collected was centrifuged at 4000 rpm for 10 min and the serum was removed and stored at −80°C to analyze for cholesterol, HDL, LDL, VLDL, triacylglycerol, total serum proteins, albumin and activities of the enzymes alanine aminotransferase (ALT) and alkaline phosphatase (ALK). All the serum biochemical analyses were performed using commercial kits (Labtest®) in an automatic serum biochemistry analyzer (Wiener lab group, Rosário-Argentina). Ca and Pi content in sera was determined using commercial colorimetric kits (Labtest®) immediately after the blood sampling.

### 2.6 Computed tomography (CT) scan and bone density

Two fish per tank were analyzed in a helicoidal CT-scanner (Shimadzu, model SCT-7800 CT) using the following parameters: 250 and 350 mm of visual field according to the fish size, algorithm 120 kv and 100 ma, section interval of 2 mm (gap) and 2 mm section thickness. The tomographic analysis of axial sections showed images of bones and vertebrae in the right side position and allowed quantitative measurements of the bone mineral density by visualization of the alterations in bone structure to compare the treatments. Images were analyzed using the software ClearCanvas Workstation®. The bone density was calculated according to the Hounsfield scale using 0.070 cm^2^ as the standard diameter. Three vertebral bodies were measured per fish in relation to the pectoral fin, dorsal fin and caudal fin, thus, nine measurements were performed in each fish (Fig. 1).

**Fig 1.**
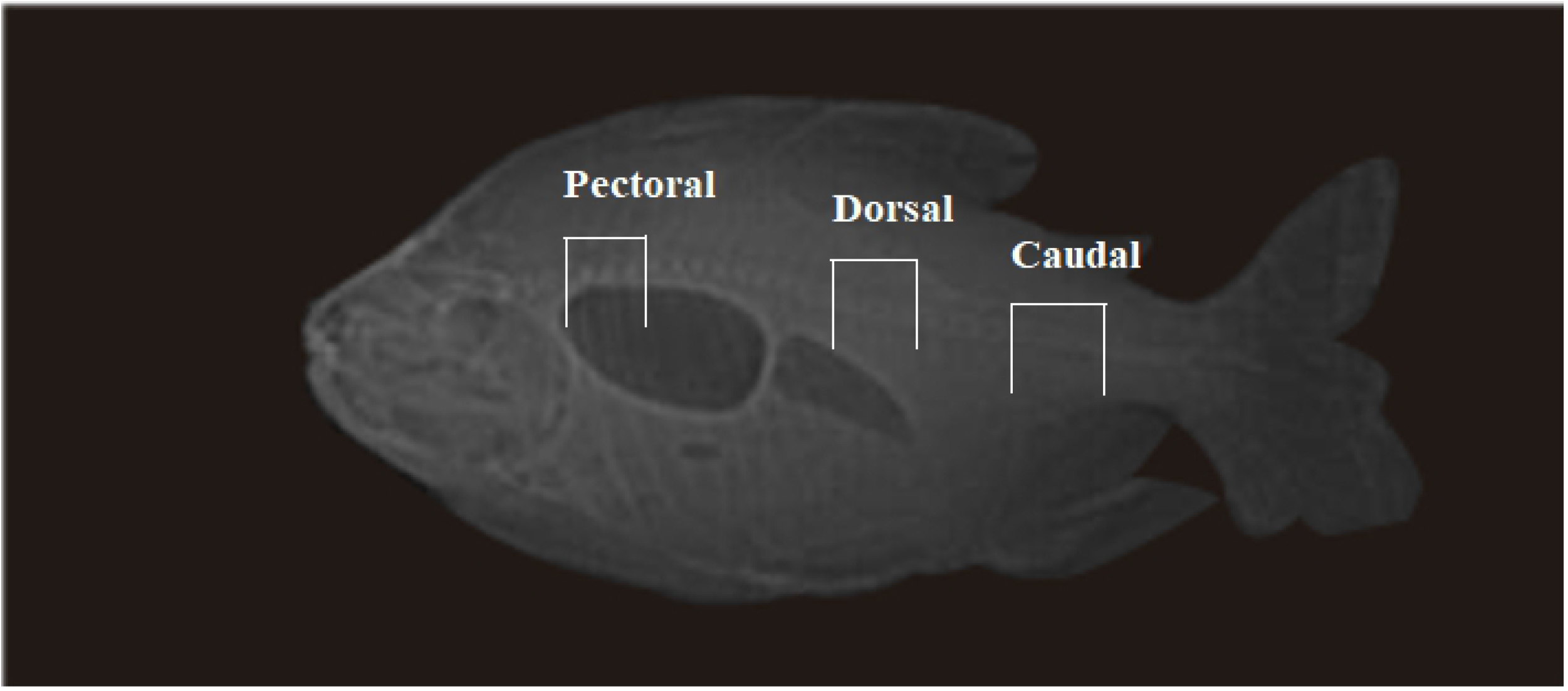
CT-scanning images of tambaqui fed diets containing graded levels of digestible phosphorus showing the theer distinct regions considered during measurement of the bone mineral density of the vertebral: Pectoral, Dorsal and caudal.

### 2.7 Statistical analysis

All data were checked for normality (Shapiro Wilk test) and homoscedasticity (Bartlett test). Then, the ANOVA procedure was applied and when significant differences were observed (P<0.05), linear and non-linear regression models were tested to check the best fit. Models were selected based on the least sum of squared differences between the values of the observed and predicted values of the dependent variable [19], the P value and the R^2^ [20]. Because we do not want to compare body weight responses to P (the fixed-effect viewpoint), rather we want to estimate the overall response of a population, requirements were estimated using all data without considering the block effect which were significant for some response parameters (Appendix S1). For the variables which the quadratic equation was the best model, the requirement was estimated based on 95% of the maximum response [19]. For response variables that were significantly different, and no model was able to be fitted, pairwise comparisons between treatments means were made using the Student–Newman–Keuls or Duncan multiple range test. All data were analyzed using the easyanova and easyreg packages of R*®* software and graphs were prepared using the GraphPad Prism software.

## 3. Results

The analyzed total phosphorus levels in the diets were similar to the calculated levels in the formulation (Table 1). Fish accepted well the semi-purified diets and showed a normal feeding behavior throughout the experiment. All growth performance parameters were affected by dietary phosphorus (P<0.05) (Table 2). The maximum growth rate was estimated to be at 6.3 dP g kg^−1^ diet. Fish fed diets containing dP levels lower than those had a significant reduction on feed intake, specific growth rate and impaired feed conversion ratio. Protein efficiency ratio and protein retention linearly increased according to the dietary dP levels. On the other hand, dietary P was better utilized by fish fed the lowest dP levels. The estimated requirement based on growth parameters varied from 3.7 to 7.1 and different models best fitted to the parameters (Table 3). No mortalities were observed throughout the experiment.

**Table 2.**
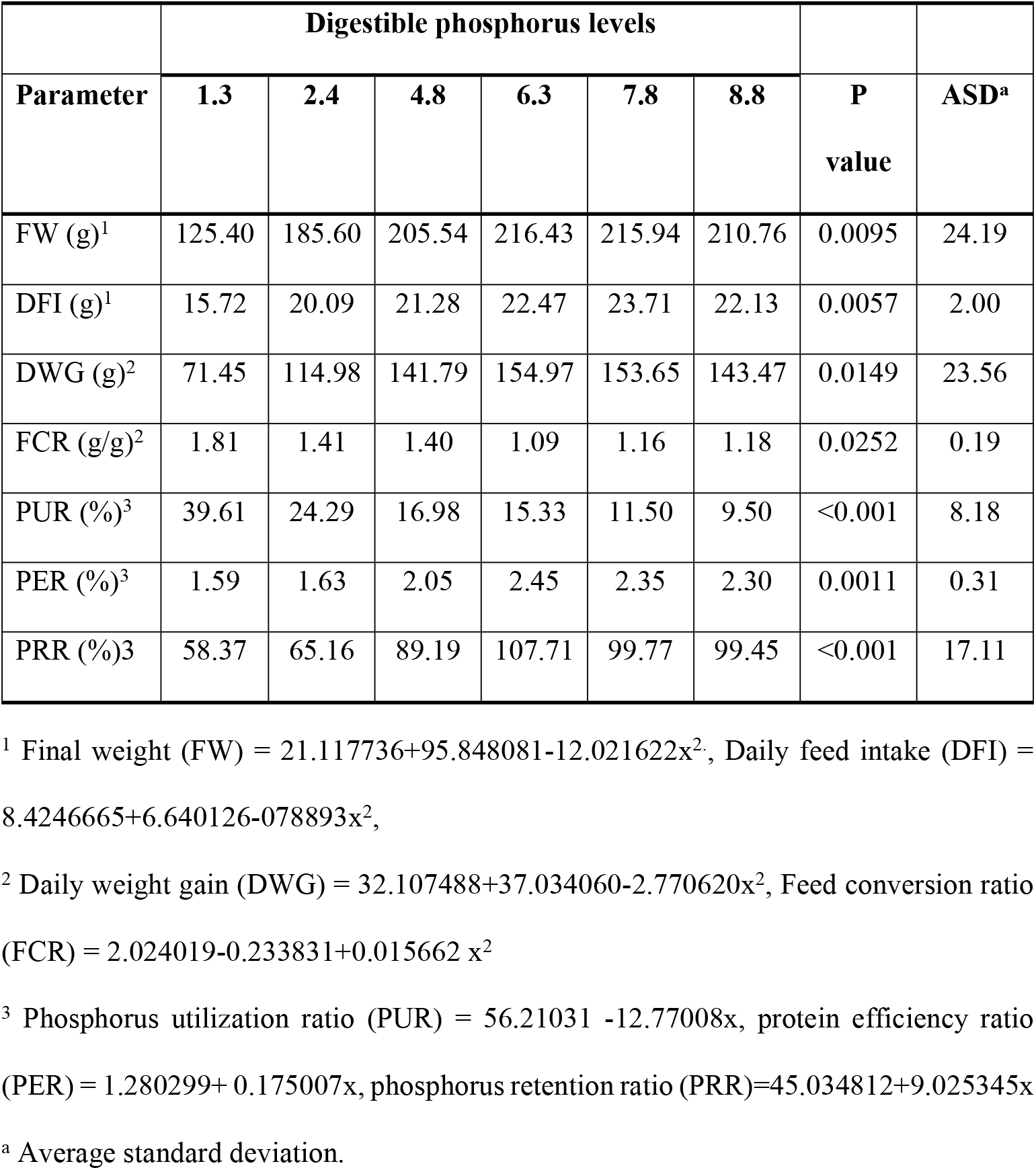
Growth performance of tambaqui fed diets containing graded levels of digestible phosphorus

**Table 3.** Best fitted regression models, P values, Mean Squared Residue (MSR) and estimated phosphorus requirement of the growth parameters of tambaqui fed diets containing graded levels of digestible phosphorus

| Variable | Regression model | R<sup>2</sup> | P value | MSR | Requirement<br>(g Kg<sup>-1</sup>) |
| --- | --- | --- | --- | --- | --- |
| FW | Quadratic plateau | 0.98 | 0.0083 | 1973.75 | 3.76 |
| DFI | Quadratic plateau | 0.92 | 0.0092 | 12.3161 | 3.96 |
| DWG | Quadratic | 0.98 | 0.0082 | 1652.10 | 6.33 |
| FCR | Quadratic | 0.77 | 0.0011 | 0.1048 | 7.10 |
| PUR | Broken line linear<br>plateau | 0.94 | <0.001 | 30.12 | 3.18 |
| PER | Broken line linear<br>plateau | 0.96 | <0.001 | 0.1084 | 5.96 |
| PRR | Broken line linear<br>plateau | 0.97 | <0.001 | 270.7408 | 6.32 |

Whole-body lipid significantly decreased while the ash content linearly increased in response to the dietary dP levels (Table 4). Regardless for Ca and P, dietary dP levels did not affect the whole-body protein and moisture content (Table 4). P content in and scales showed a quadratic effect in response to dP supplementation (Table 6). Estimated requirement based on carcass gross nutrient composition was around 6.6g kg^−1^ diet, while for vertebrae mineralization was between 4.75 and 6.75 g kg^−1^ diet (Table 5 and 6). However, mineral deposition on scales varied greatly between phosphorus and calcium (Ca). Estimated dP requirement based on scales P deposition was 7.38 g kg^−1^ while the estimated requirement for Ca deposition was 4.04 g kg^−1^.

**Table 4.** Whole-body chemical composition of tambaqui fed diets containing graded digestible phosphorus levels (n = 8, dry matter basis)

|  | **Digestible phosphorus levels** |  |  |  |  |  |  |  |
| --- | --- | --- | --- | --- | --- | --- | --- | --- |
| **Variable** | **1.3** | **2.4** | **4.8** | **6.3** | **7.8** | **8.8** | **P value** | **ASD<sup>a</sup>** |
| Moisture | 55.49 | 57.13 | 60.17 | 59.95 | 57.96 | 58.15 | 0.1251 | 1.28 |
| Crude protein | 45.70 | 47.67 | 48.63 | 47.93 | 46.95 | 48.35 | 0.4211 | 0.81 |
| Lipid<sup>1</sup> | 39.89 | 39.99 | 37.71 | 33.24 | 35.11 | 33.99 | 0.0585 | 2.54 |
| Ash<sup>1</sup> | 6.60 | 7.27 | 8.82 | 11.21 | 10.16 | 12.21 | 0.0433 | 1.81 |

<sup>1</sup> Lipid = 41.36686 - 0.900377x, Ash = 5.679903 + 0.70663x
<sup>a</sup> Average standard deviation.

**Table 5.** Mineral composition of whole-body, bone vertebrae, and scales of tambaqui fed diets containing graded levels of digestible phosphorus levels (n=8, g kg^−1^ for Ca, P and Mg; mg kg^−1^ for Mn and Zn)

|  | Digestible phosphorus levels (g kg<sup>-1</sup>) |  |  |  |  |  |  |
| --- | --- | --- | --- | --- | --- | --- | --- |
| Variables | 1.3 | 2.4 | 4.8 | 6.3 | 7.8 | 8.8 | ASD<sup>a</sup> |
| **Whole-body** |  |  |  |  |  |  |  |
| Phosphorus (g Kg<sup>-1</sup>)<sup>1</sup> | 29.17 | 39.19 | 46.37 | 50.53 | 52.39 | 48.02 | 0.73 |
| Calcium (g Kg<sup>-1</sup>)<sup>2</sup> | 59.85 | 60.84 | 67.21 | 62.50 | 72.57 | 72.36 | 0.48 |
| Magnesium (g Kg<sup>-1</sup>) | 0.17 | 0.17 | 0.18 | 0.16 | 0.15 | 0.14 | 0.002 |
| Zinc (mg Kg<sup>-1</sup>) | 1135 | 1471 | 1158 | 1361 | 1311 | 1367 | 0.012 |
| **Vertebrae** |  |  |  |  |  |  |  |
| Phosphorus (g Kg<sup>-1</sup>)<sup>1</sup> | 130.94 | 158.81 | 171.14 | 173.27 | 169.83 | 178.63 | 1.25 |
| Calcium (g Kg<sup>-1</sup>)<sup>2</sup> | 143.15 | 163.64 | 196.64 | 208.33 | 226.74 | 193.19 | 4.42 |
| Magnesium (g Kg<sup>-1</sup>) | 3.02 | 6.06 | 3.95 | 2.65 | 3.29 | 2.24 | 0.06 |
| Manganese (mg Kg<sup>-1</sup>) | 90 | 91 | 89 | 87 | 92 | ND | 0.00014 |
| Zinc (mg Kg<sup>-1</sup>) | 157 | 121 | 147 | 145 | 143 | ND | 0.00086 |
| **Scales** |  |  |  |  |  |  |  |
| Phosphorus (g Kg<sup>-1</sup>)<sup>1</sup> | 63.10 | 76.83 | 87.65 | 96.07 | 97.67 | 95.59 | 1.07 |
| Calcium (g Kg<sup>-1</sup>)<sup>3</sup> | 49.38 | 55.04 | 60.10 | 66.78 | 83.87 | 90.03 | 1.29 |
| Magnesium (g Kg<sup>-1</sup>) | 2.90 | 3.69 | 3.79 | 3.53 | 3.01 | 4.75 | 0.04 |
| Zinc (mg Kg<sup>-1</sup>) | 387 | 358 | 359.00 | 332 | 310 | 357 | 0.00197 |

<sup>1</sup> Whole-body phosphorus = $1.939912 + 0.904205x - 0.064585x^2$ , Scale phosphorus = $4.996283 + 1.204647x - 0.077468x^2$ , Vertebrae phosphorus = $12.546399 + 3.040429x - 0.303498x^2$ , Vertebrae calcium = $10.652064 + 2.961177x - 0.208294x^2$
<sup>2</sup> Whole-body calcium = $5.89803270 + 0.103855x$
<sup>3</sup> Scale calcium = $2.018257 + 0.792441x$
<sup>a</sup> Average standard deviation.
Bars with different letters are significantly different by SNK multiple range test ( $P < 0.05$ ).

**Table 6.**
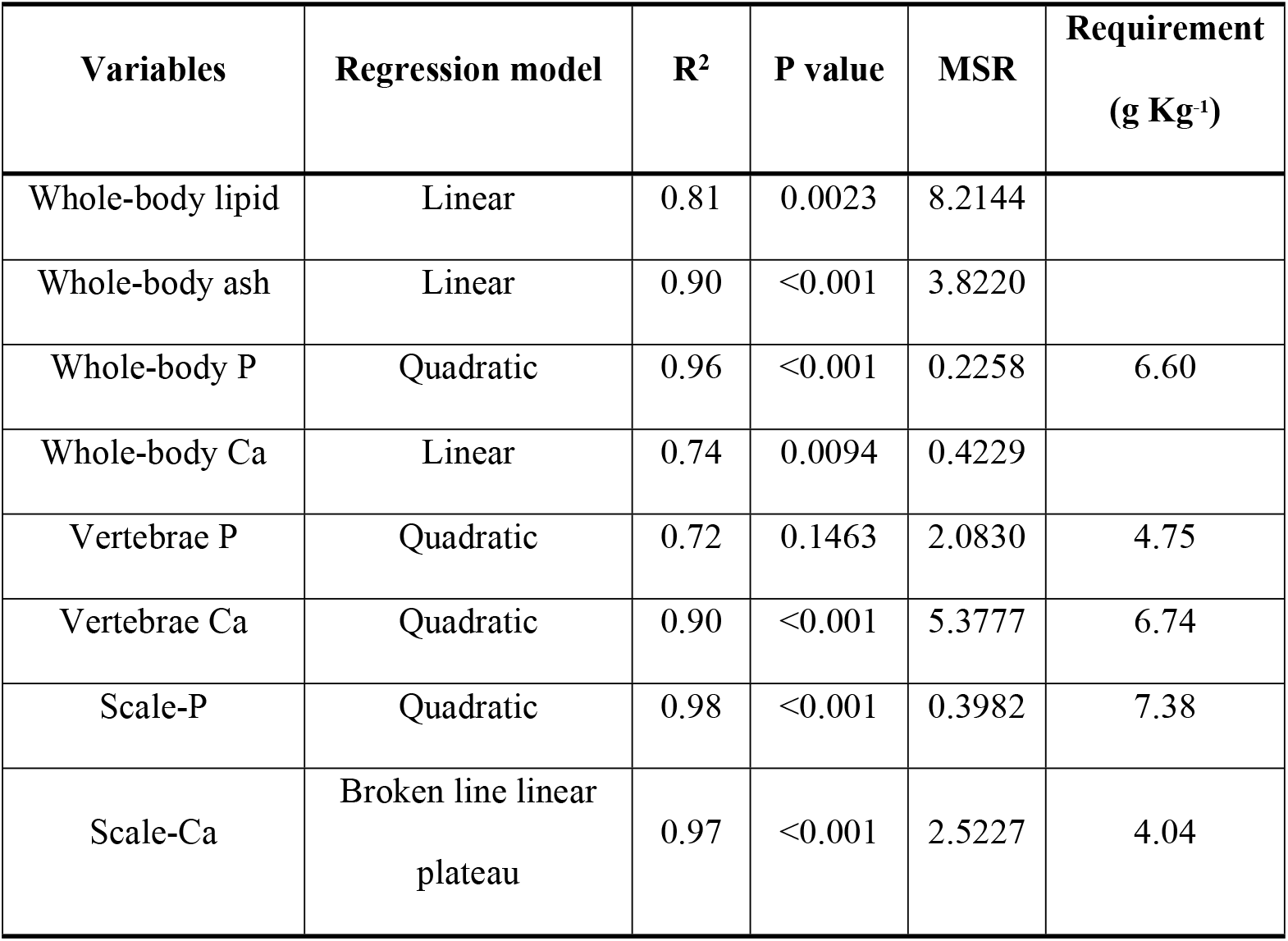
Regression model, P value, Mean Squared Residues (MSR), and P requirement of chemical composition of body compartments and bone mineralization of tambaqui fed diets containing graded levels of digestible phosphorus

| Variables | Regression model | R<sup>2</sup> | P value | MSR | Requirement (g Kg<sup>-1</sup>) |
| --- | --- | --- | --- | --- | --- |
| Whole-body lipid | Linear | 0.81 | 0.0023 | 8.2144 |  |
| Whole-body ash | Linear | 0.90 | <0.001 | 3.8220 |  |
| Whole-body P | Quadratic | 0.96 | <0.001 | 0.2258 | 6.60 |
| Whole-body Ca | Linear | 0.74 | 0.0094 | 0.4229 |  |
| Vertebrae P | Quadratic | 0.72 | 0.1463 | 2.0830 | 4.75 |
| Vertebrae Ca | Quadratic | 0.90 | <0.001 | 5.3777 | 6.74 |
| Scale-P | Quadratic | 0.98 | <0.001 | 0.3982 | 7.38 |
| Scale-Ca | Broken line linear plateau | 0.97 | <0.001 | 2.5227 | 4.04 |

Bone density of tambaqui fed diets containing graded levels of dP quadratically increased with an estimated requirement around 3.68 g kg^−1^ and then plateau. The exception of the set of caudal vertebrates that linearly increased up to 2.76 g kg^−1^ and then plateau. The density of the vertebral bodies in tambaqui was approximately 20% higher in the pectoral vertebrae, except for the fish fed the basal diet which had similar densities in all vertebral regions (Table 7).

**Table 7.** Bone density of tambaqui fed diets containing graded levels of digestible phosphorus (n=8)

|  | Dietary digestible phosphorus levels |  |  |  |  |  |  |
| --- | --- | --- | --- | --- | --- | --- | --- |
| Vertebrae | 1.3 | 2.4 | 4.8 | 6.3 | 7.8 | 8.8 | ASD<sup>a</sup> |
| Pectoral (HU)<sup>1</sup> | 297.7 | 412.95 | 429.85 | 464.05 | 482.94 | 463.19 | 46.5256 |
| Dorsal (HU)<sup>1</sup> | 261.07 | 334.3 | 340.71 | 366.82 | 383.15 | 353.56 | 28.1667 |
| Caudal (HU)<sup>2</sup> | 268.32 | 346.9 | 345.43 | 373.42 | 408.45 | 365.53 | 31.125 |

HU: Hounsfield unit
<sup>1</sup> Quadratic plateau effect equations: Pectoral = $72.34042 + 210.5015x - 28.57533x^2$ ( $R^2=0.93$ ), Dorsal = $114.4589 + 137.8055x - 19.2521x^2$ ( $R^2= 0.89$ ),
<sup>2</sup> Broken line linear plateau effect: Caudal = $175.4527 + 71.43636x$ ( $R^2=0.81$ )
<sup>a</sup> Average standard deviation.

Serum Pi levels were highest in fish fed diets supplemented with 4.8, 6.3 and 7.8 g kg^−1^ dP, while the lowest values were recorded for fish fed the two lowest dP levels (Fig. 2A). Serum albumin concentration linearly increased up to 3.18 g kg^−1^ and then plateau (Fig. 3B), while the plateau value for ALB:GLOB ratio was 3.92 g kg^−1^ according to the increase on dietary dP levels. Serum lipid profile of tambaqui was profoundly affected by the dietary dP levels (Fig. 4). Generally, TAG and VLDL were higher in fish fed the three lowest dP levels (Fig 4A and 4E, respectively), while LDL showed a linear increase with a plateau at 4.76 (Fig. 4D). The highest total cholesterol level was recorded in fish fed diets with the two highest dP levels (Fig. 1B). Similarly, the enzyme activity was affected by dietary dP levels (Fig. 5). ALK activity linearly decreased up to 3.08 g kg^−1^ and then plateau in response to dietary dP levels (Fig. 5A).

**Fig 2.**
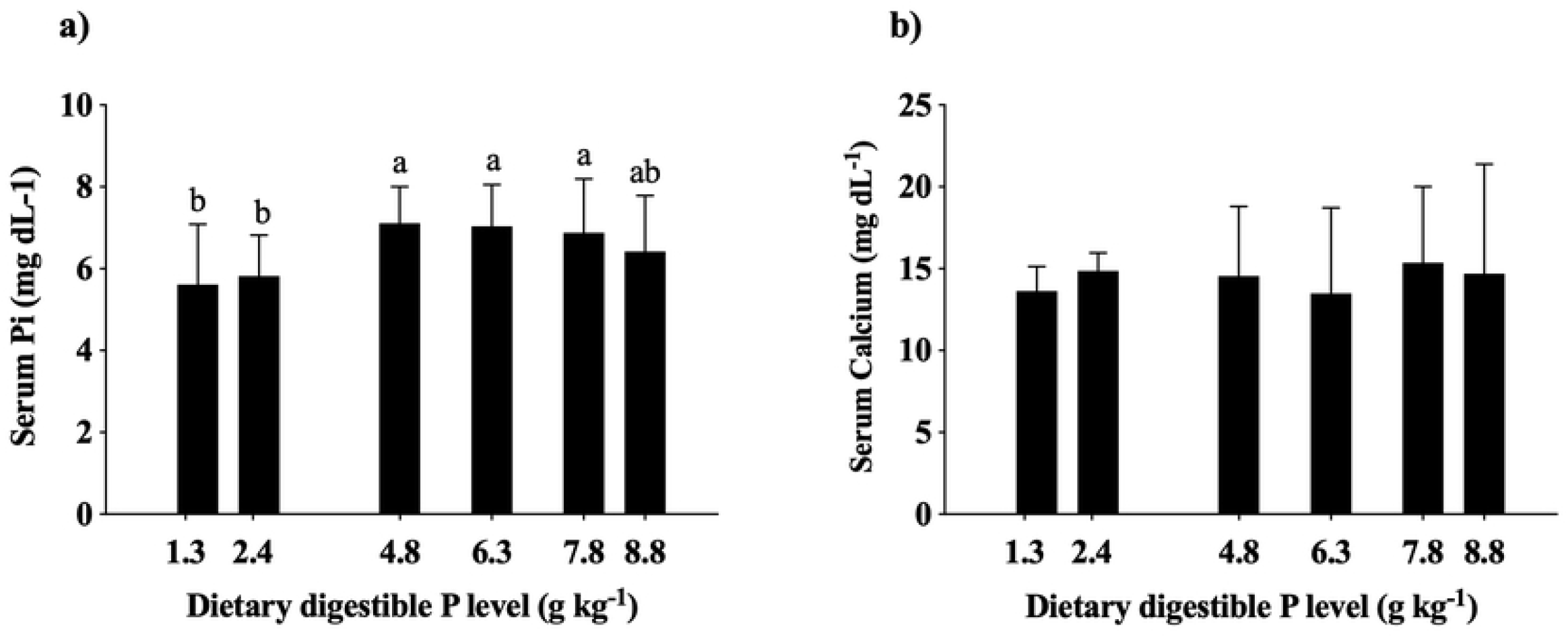
Serum phosphorus (A) and calcium (B) concentration of tambaqui fed diets containing graded levels of digestible phosphorus (n=12) Bars with different letters are significantly different by Duncan multiple range test (P < 0.05).

**Fig 3.**
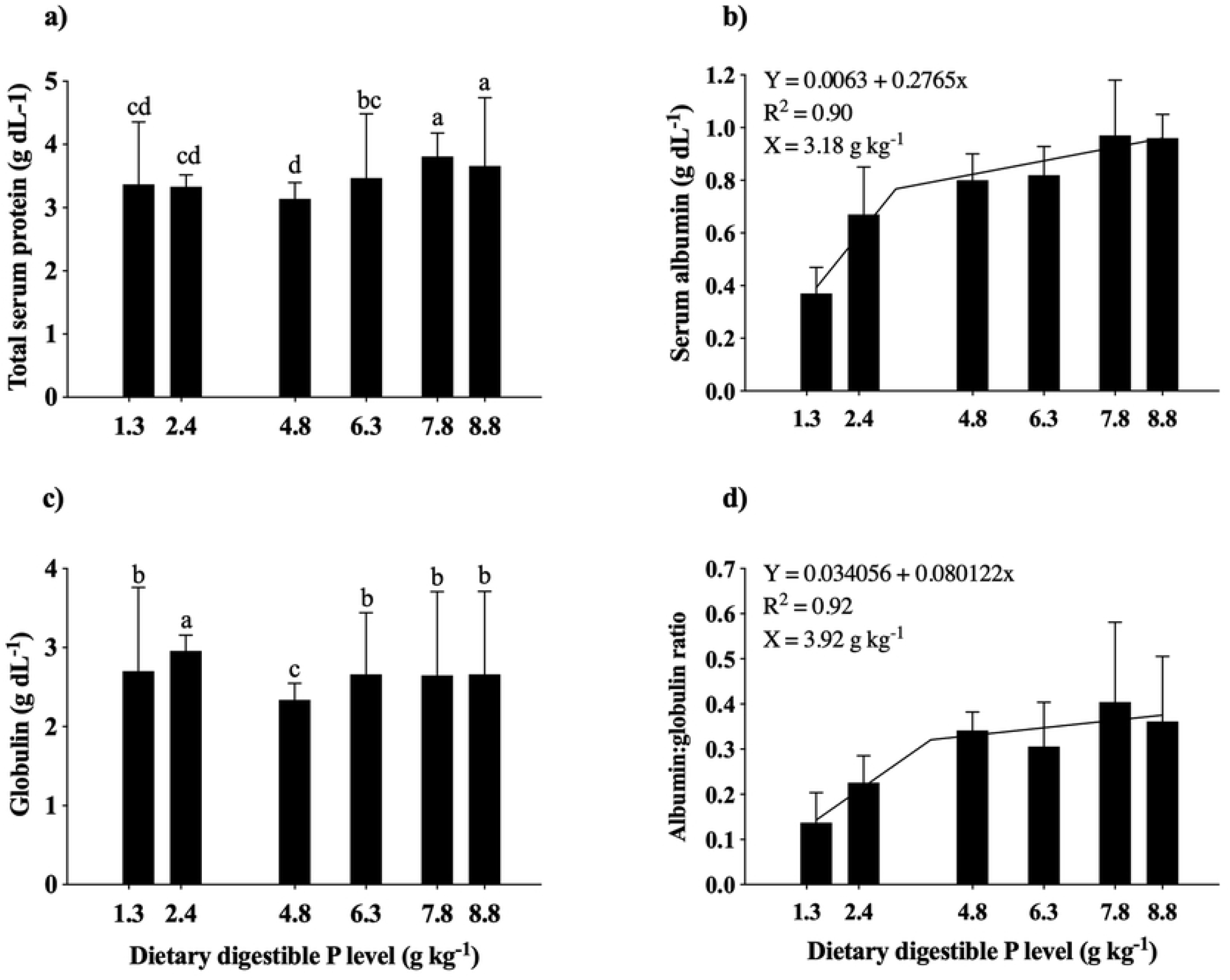
Protein (A), albumin (B), globulin (C) concentration and ALB:GLOB ratio (D) in tambaqui sera fed diets containing graded levels of digestible phosphorus (n=12). Bars with different letters are significantly different by Duncan multiple range test (P < 0.05); ^1^Broken line linear plateau regression

**Fig 4.**
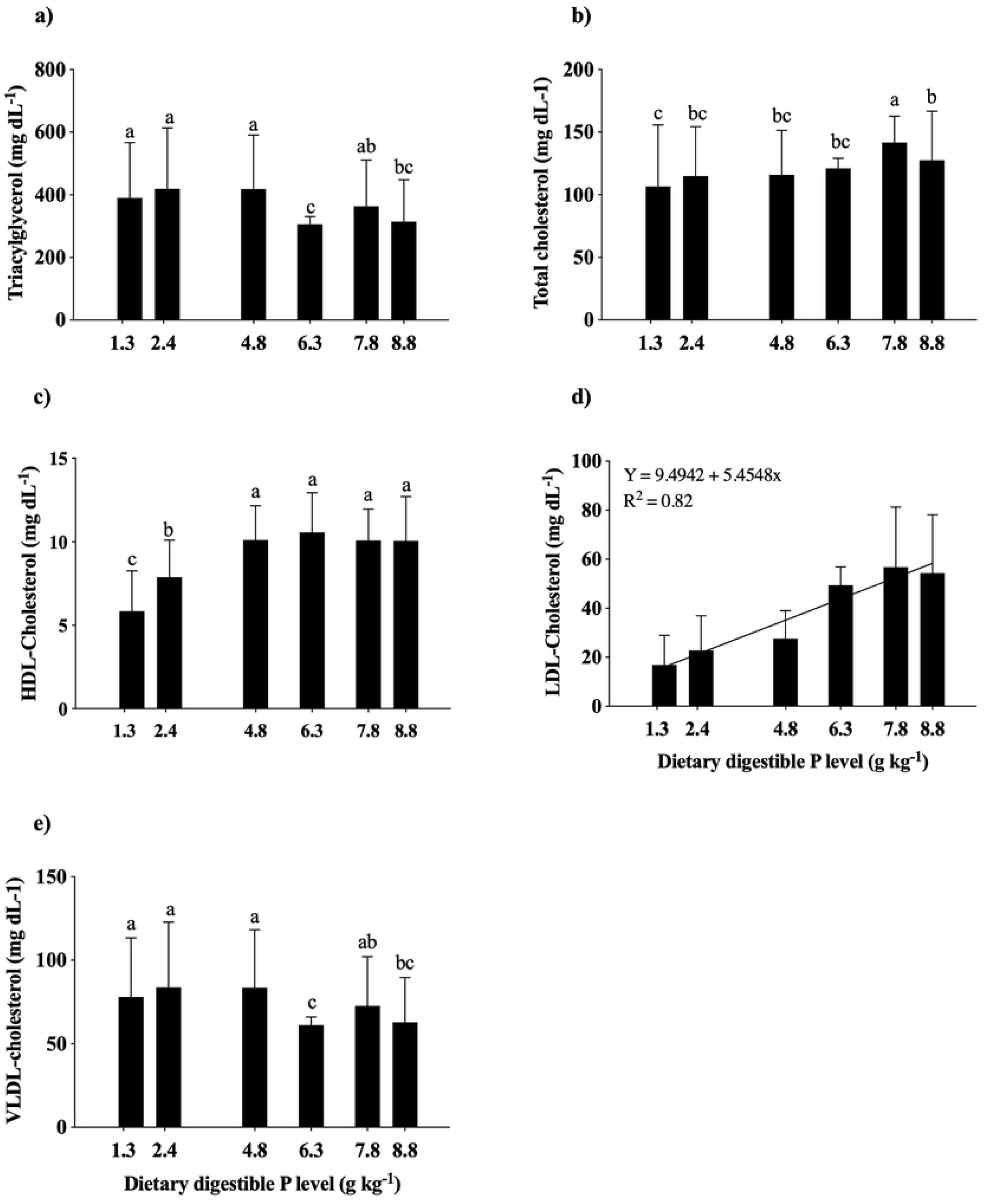
Triacilglycerol (A), cholesterol (B), HDL-cholesterol (C), LDL-cholesterol (D) and VLDL (E) concentration in tambaqui sera fed diets containing graded levels of digestible phosphorus (n=12). Bars with different letters are significantly different by Duncan multiple range test (P < 0.05); ^1^Broken line linear plateau regression

**Fig 5.**
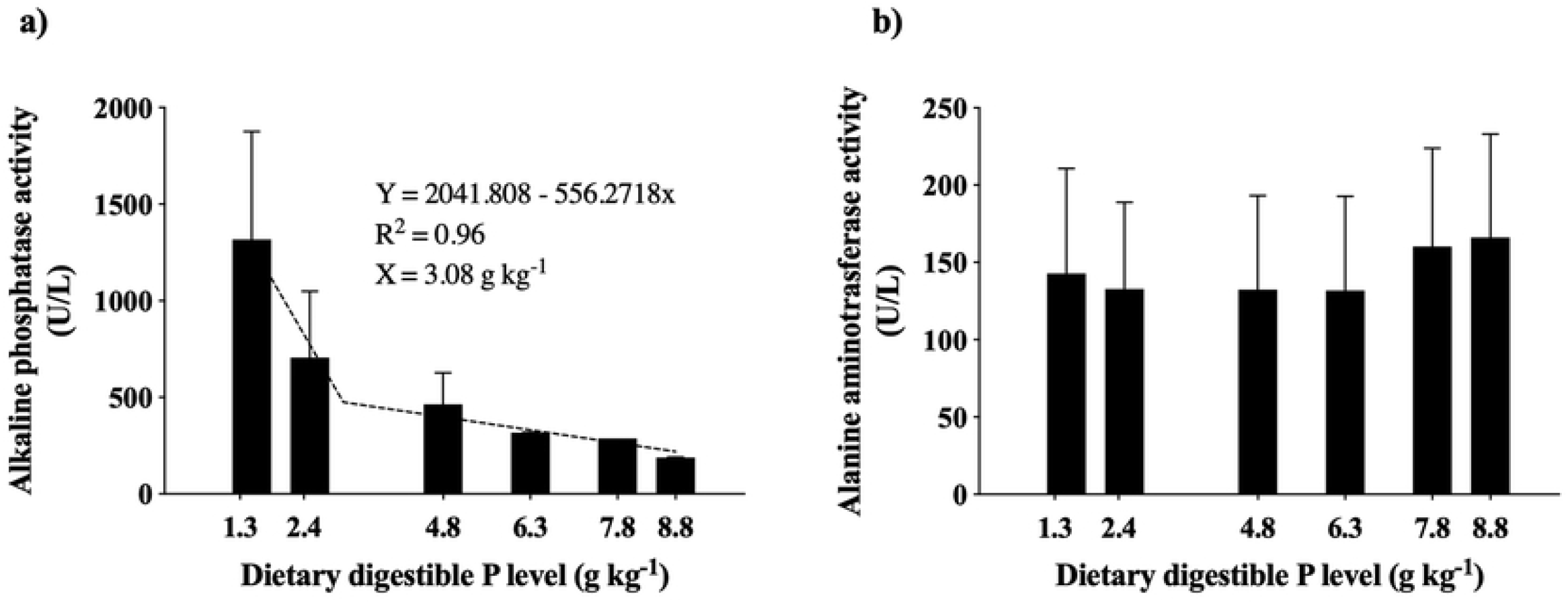
Serum alkaline phosphatase (A) and alanine aminotransferase (B) activities of tambaqui fed diets containing graded levels of digestible phosphorus (n=12). Bars with different letters are significantly different by Duncan multiple range test (P < 0.05); ^1^Broken line linear plateau regression

## 4. Discussion

Phosphorus deficiency in tambaqui was characterized by retarded growth rate and impairment of all growth parameters. Indeed, the main symptoms of phosphorus deficiency in fish are slower growth rates, lower feed intake and reduced nutrient utilization [21]. However, the efficiency of phosphorus utilization was higher in fish fed P-deficient diets. Similar results have been previously reported with different species, such as *Takifugu obscurus* [22], *Cirinius carpio* [23, 24], *Hucho taimen* [10], *Oreochomis niloticus* [25], *Carassius auratus gibelio* [26], *Carassius auratus[27], Sparus macrocephalus[9]*. This effect is usually related to a physiological adaptation of the gut and kidney cells to improve P uptake by increasing the expression of NAPi^+^ transporters, thus, increasing P absorption and reducing P excretion [28].

Tambaqui juveniles reached the maximum growth rate with 6.3 g kg^−1^ diet of dP. This value is higher than a previous study with tambaqui that reported a dP requirement of 3 g kg^−1^ diet at the beginning of the grow-out phase (150 to 300g) [7]. Since, there was a great difference between the requirement estimates for tambaqui, we believe the previously reported requirement might be underestimated once they used plant based practical diets and there are evidences showing that tambaqui might be able to efficiently use dietary phytic-P [15]. Although the P requirement tend to reduce according to the increase of growth stage [5], our estimated P requirement was extremely higher than the previous study.

Mineralization of bones, scales and whole-body was reduced in fish fed P-deficient diets. This well-characterized P deficiency symptom in fish [29] [30] [9] occur due to an intensive mobilization of calcium phosphates of mineralized structures to prioritize physiological processes [21]. This hypothesis corroborates with the high ALK activity in fish fed P-deficient diets. In higher vertebrates, ALK activity is used as a biomarker of bone turnover since the cells responsible for bone remodeling, osteoblasts, secrete high amount of ALK [31]. Additionally, the scales are an important mineral storage compartment in fish and are highly responsive to P deficiency. Recent studies indicated that besides scales and bones share similar metabolic pathways, mineral mobilization occurs preferably through scales in P deficiency in order to maintain bone structure [11]. Thus, the P levels in scales might be a useful non-invasive tool to estimate mineral requirement in fish. The requirement for proper scale mineralization in this study was slightly higher (7.38 g kg^−1^) than the growth estimation and whole-body mineralization. This indicates that scales might have an important function on maintaining mineral homeostasis in fish.

P requirement based on bone mineralization are generally higher than the values estimated based on growth or whole-body mineralization [21, 29, 32]. However, our results with tambaqui surprisingly showed the contrary (4.75 versus 6.33 g kg^−1^ for bone mineralization and growth, respectively). We hypothesize that the P requirement based on growth is greatly affected by the amount of mineralized structures (such as scales, operculum and fins) of the species. Comparing tambaqui with Atlantic salmon, one of the most studied species in aquaculture, tambaqui seems to have a higher number of mineralized structures and bone density than salmonids and therefore this might affect the determination of the requirement. In fact, this hypothesis was supported by our findings that fish fed the deficient diet clearly showed a reduced number of mineralized structures and the total bone volume was 2.90 cm^3^, while fish fed adequate P levels (6.3 g Kg-^1^) showed a great increase on total bone volume (13.34 cm^3^). Thus, the assumption that the requirement based on the bone mineralization is higher than those based on growth might not be true for all fish species. Additionally, the CT-scanning could predict the P requirement for bone mineralization using the vertebrae as the bone tissue.

In this view, the evaluation of bone density could be an important tool to assess and/or properly interpret the data on P requirement studies, since bone mineralization is reduced by P deficiency with a concurrent reduction on bone density [33]. However, few studies have used this technique to evaluate P adequacy in fish and, to the best of our knowledge, this is the first study to report the use of CT-scan to determine the mineral requirement of a fish species. In general, most of the studies use a qualitative measure obtained by X-ray analysis. The evaluation using the X-ray method is described for a fewer fish species, such as Atlantic salmon, [33], *Senegalese sole* [30], *Melanogrammus aeglefinus* L. [18] and rainbow trout [12] fed graded levels of P. In this study, we report the bone density based on CT-scan evaluation. This method converts the shades of grays of the X-ray analysis on numeric values, called Housenfield units [33, 34] allowing us to have an isolated measure of the fish vertebrae density. Although we could not observe any macroscopic lesions on deficient fish, the 3D representation of the CT-scan (Fig. 6) allowed the visualization of impaired bone mineralization in several body structures of the fish. Therefore, we could see that the main bone structures affected by P deficiency are the cranial and opercular bones.

**Fig 6.**
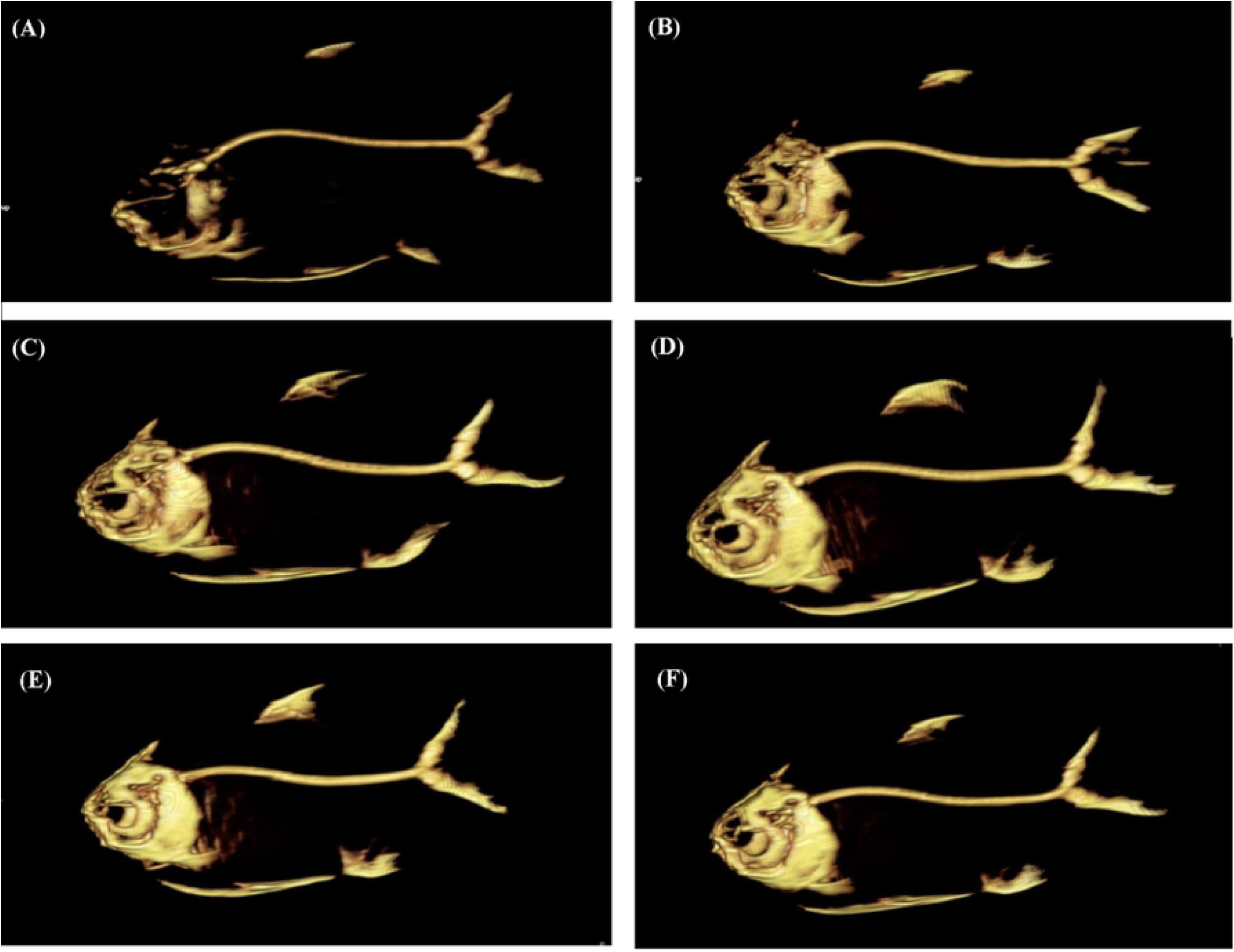
CT-scanning images of mineralized tissues in tambaqui fed diets containing graded levels of digestible phosphorus; Fishes fed basal diet (A), 2.5 g kg-1 dP diet (B), 4.8 g kg-1 dP diet (C), 6.3 g kg-1 dP diet (D), 7.8 g kg-1 dP diet (E), and 8.8 g kg-1 dP diet (F).

The increase on fatty body deposition is a commonly reported effect of P deficiency in fish [4, 7, 10, 13, 24–26, 30, 35]. In this study, we have similarly observed an increase on body fat content with a concomitant increase on triacylglycerol and VLDL levels, and a decrease on serum albumin levels. Although this effect is not completely elucidated in fish, there are compelling evidences that this increase on fat deposition is a combining effect of the inhibition of the β-oxidation and the use of amino acids as an energy supply as this can be supported by the serum biochemical results in our study. Additionally, a previous study on carp have observed an increase on activity of citrate lyase, a rate limiting enzyme of the lipogenesis, and the gluconeogenesis enzymes: GDH (glutamate dehydrogenase), PEPCK (phosphoenol pyruvate kinase) and FBP (fructose 1,6-biphosphatase) [36]. The authors already observed a reduction on pyruvate kinase activity, a rate limiting enzyme for the use of glucose as an energy source. Recently, a study with *Takifugu obscurus* showed that dietary P positively upregulated the enzymes of pentose phosphate shunt, which are the main NADPH supply for the lipogenesis [22]. Altogether, this reinforces the hypothesis that P deficient fish obtain the energy mainly through the amino acid/ protein catabolism while the glucose is being used as the main substrate for lipogenesis through the pentose phosphate shunt.

The ammonia excretion is the main biomarker for the protein catabolism in fish and in P-deficient fish there is an increased post-prandial ammonia excretion as have been reported for *Bidyanus bidyanus* [29, 37]. These findings are in one hand with the reduced protein efficiency and retention in P-deficient fish of this study. Furthermore, other studies have reported similar results with other fish species, such as *Carassius auratus* [27], *Oreochromis niloticus* [8], *Senegalese sole* [30], *Carassius auratus gibelio* [26].

The study of the blood components is an important tool to evaluate the welfare and overall physiological conditions. For instance, the cholesterol plays an important role on maintaining proper membrane permeability and fluidity being considered an adequate biomarker of fish welfare [38]. In general, fish has higher cholesterol levels compared to higher vertebrates. Therefore, lower cholesterol levels might be an indication of physiological distress [38–40]. In this study, P-deficient fish showed a lower total and LDL cholesterol levels. This reduced level of cholesterol might be an effect of the use of cholesterol in the synthesis of corticosteroids. However, few studies have evaluated the effect of P supplementation on cholesterol metabolism in fish and the results are contradictory. For instance, P-deficiency in *Takifugu obscurus* [22] induced lower serum cholesterol levels while the contrary was observed in *Sparus macrocephalus* [9] e *Ctenopharyngodon idella* [23]. Therefore, this effect of P on cholesterol levels should be further studied in fish since the mechanism used to explain its action is based on higher vertebrates which might not be similar. Additionally, this might be an effect of the high fat deposition in the liver in P-deficient fish. The low serum ALB:GLOB ratio and higher ALT activity might be an indication of liver damage and concurrent inflammation since these parameters are usually used as biomarkers to assess the health status in higher vertebrates [41].

The use of serum Pi as a biomarker to assess the proper dietary P levels in fish has been criticized by several studies because the serum P usually reflects the effect of a recent meal rather than P status of a fish [7, 35, 42]. However, we found serum Pi to be a responsive biomarker of P status in this study since its results correlates well with all the other parameters used to assess the dietary P adequacy. This effect is supported by the serum albumin levels, which showed the same trend as the Pi levels. Albumin is responsible for the transport of several molecules through the body, phosphate included [7]. Therefore, in P-deficient animals the low Pi levels are usually followed by reduced albumin levels. This effect might have been prominent in our study because the fish were fasted for 24h before the blood sampling, thus reducing the effect of the last meal.

Taking altogether, our study demonstrated that feeding tambaqui with 6.3 g kg^−1^ digestible P on the beginning of the growth stage is adequate based on several parameters of growth and welfare. Additionally, we have demonstrated for the first time that CT-scan might be an adequate and non-invasive method to assess the P requirement or bone health of fish. Nonetheless, the use of vertebrae mineralization as a response parameter might not be adequate for the welfare of some fish species. Further studies on the effect of P status on lipid metabolism and other blood chemistry parameters are warranted since a profound effect of P status on these parameters was observed in this study.

## 5. Acknowledgements

This research was funded by Fundação de Amparo a Pesquisa do Estado de Goiás - FAPEG (grant no 201410267000859). We are thankful to Guabi Nutrição e Saúde Animal S.A and CJ Selecta for providing the micronutrient mix and the soy protein concentrate, respectively. We also are thankful to PUC-GO for providing their research facilities to conduct the trial. Janaína G. Araújo post doc and Ludmila L.C. Menezes Master scholarships were supported by CNPq and Coordination of the Improvement of Higher Level Education (CAPES, Brazil), respectively.

